# Enhanced Plasmid-Based Transcriptional Activation in Developing Mouse Photoreceptors

**DOI:** 10.1101/2024.06.06.597220

**Authors:** Brendon M. Patierno, Mark M. Emerson

## Abstract

Rod photoreceptor formation in the postnatal mouse is a widely used model system for studying mammalian photoreceptor development. This experimental paradigm provides opportunities for both gain and loss-of-function studies which can be accomplished through in vivo plasmid delivery and electroporation. However, the cis-regulatory elements used to implement this approach have not been fully evaluated or optimized for the unique transcriptional environment of photoreceptors. Here we report that the use of a photoreceptor cis-regulatory element from the Crx gene in combination with broadly active promoter elements can increase the targeting of developing rod photoreceptors in the mouse. This can lead to greater reporter expression, as well as enhanced misexpression and loss-of-function phenotypes in these cells. This study also highlights the importance of identifying and testing relevant cis-regulatory elements when planning cell subtype specific experiments. The use of the specific hybrid elements in this study will provide a more efficacious gene delivery system to study mammalian photoreceptor formation.

## Introduction

The vertebrate retina is a complex tissue that contains many cell types which must function together in precise organization. During development these cell types must differentiate after formation from retinal progenitor cell (RPC) division and orient within the retina appropriately. Differentially open chromatin regions of genomic DNA and cell type specific transcription factor expression within these cells define which gene regulatory networks are active in post-mitotic cells as opposed to RPCs, and differences exist even within cell subtypes. Thus, knowledge of cell type specific transcription factors and their corresponding cis-regulatory elements (CREs) is critical to studying the development of the retina.

For studies that implement gain-of-function or loss-of-function experiments during retina development, it is critically important to understand which cell types are targeted by the transcriptional sequences present in the introduced vectors. This is especially true in complex tissue types such as the retina, where many cell types interact in networks with other cells. Differences in the promoter activity of the CAG, EF1a, and CMV promoters in retina cells was first observed in rat tissue with the observation that CAG was the most active element in photoreceptors^1^. Subsequent studies in the mature mouse retina thus have most often used CAG as a ubiquitous promoter, and qualitatively observed CAG::GFP to have similar activity in vivo to what was seen in the rat retina^2^. However, upon examination of postnatal retina experiments that use the CAG element, we observed that CAG-driven reporter activity in newly differentiating photoreceptors is often weak. Indeed, the effectiveness of CAG to induce alterations in gene expression in developing photoreceptors has not been systematically tested. This can be most clearly observed when CAG plasmids are co-electroporated with photoreceptor active elements that allow for the identification of targeted photoreceptors at early postnatal timepoints. We have observed this consistently but highlight this phenomenon for the first time here using a CRE near the Crx gene that is active in photoreceptors.

The Crx gene is expressed during both cone and rod genesis, and it’s expression in mouse and chick retina tissue has been characterized as highly expressed in photoreceptor cells^3–5^. Otx2 has been suggested to be a direct regulator of Crx, and in an Otx2 conditional knockout under the control of the Crx promoter, Crx expression in photoreceptors is completely downregulated^6^. Crx temporal expression in photoreceptors has been shown to closely follow Otx2 expression with only slight delay^7^. Furthermore, birth dating experiments show that Otx2 expression is first observed in RPCs and overlaps partially with S phase labeling by tritiated thymidine^7^. Crx expression however does not overlap at all with S phase by either tritiated thymidine or propidium iodide labeling ^7,8^. These data strongly suggest that Crx is one of the earliest markers of a newborn postmitotic photoreceptor cell. Crx is therefore an excellent candidate gene for identifying CREs that may be active in these cells.

Crx expression is however also seen to a lesser degree in bipolar cells^8^, underscoring the importance of identifying the CREs that regulate this gene to determine which elements are most active in which cell types. The specific CRE further characterized in this study, CrxE1, was first identified and predicted to be photoreceptor specific based on a high number of Nrl, Nr2e3, and Crx binding sites^9^. Testing of a reporter construct using the potential enhancer sequence revealed strong activity in photoreceptors with a high degree of specificity. This element was further characterized in both chick and mouse retinal tissue, and while still mostly active in photoreceptors, it was also observed in a population of bipolar cells by ex vivo assay^10^. However, the activity of this element has not yet been characterized in vivo in mouse tissue. Taking this information together, CrxE1 may be one of the earliest CREs active in photoreceptor specific cell formation and requires further characterization.

Although changes in photoreceptor cell proportions and opsin expression phenotypes have been observed through various gain and loss-of-function experiments, it has not been explored whether these approaches could be optimized by the use of alternative or additional enhancer elements. The observed phenotype generated by using plasmid-based overexpression of Onecut1 to induce cone photoreceptor gene expression programs in postnatal mice is of particular interest^11^. Similarly, the knock down of Nrl at p0 using Cas9 based vectors has been shown to drive the induction of the short-wave-sensitive opsin protein associated with blue light transducing cones^12^. While the genes responsible have been the subject of thorough characterization, there is little known in the field regarding how photoreceptor specific CREs could be used to enhance the degree to which these phenotypes are observed. This could be important for the generation of cone cells in other potentially clinically relevant models such as human retinal organoids or induced pluripotent stem cells. Therapeutic strategies which rely on transforming retinal cells before transplantation would benefit from an increase in the number of cells which are efficiently transformed.

This study characterizes CrxE1 activity in the mouse postnatal retina in vivo for the first time. Additionally, this study explores the method of combining cell type specific enhancers with broadly active promoters in plasmid delivery systems. The observation that the CrxE1 element is strongly active in cells where CAG is weakly active or not active at all, led to the hypothesis that there is potential to increase the amount of cells which can be targeted by ubiquitous promoter based systems, specifically to include more developing photoreceptors. Currently, these experiments are frequently performed using plasmids which contain only a promoter such as CAG or EF1a. The results of this study demonstrate the critical nature of choosing cell type specific DNA elements when performing gain or loss-of-function experiments. Specifically, the CrxE1 element represents a powerful tool to target developing photoreceptors in the post-natal mouse.

## Methods

### Animals

All procedures involving animals were approved and conducted in accordance with the City College of New York Institutional Animal Care and Use Committee and with IACUC approved protocols. CD-1 mice were obtained from Charles River.

### Molecular Biology

CrxE1::GFP^10^ (referred to as CrxEnh1 in ref. 10), which was derived from the sequence identified by Hsiau et al^9^, CAG::TdT^13^, CAG::mCherry^14^, CAG::OC1^11^, as well as empty vectors including Stagia^15^ and px458^16^ have been previously described. The CrxE1/CAG::GFP element was made using cloning strategies described here within (Figure 2). The CrxE1/CAG::OC1 element was cloned using polymerase chain reaction (PCR) with the high fidelity Herculase II Fusion DNA Polymerase (Agilent cat# 600675) to add “Sal1” sites to the 261bp enhancer sequence, and then using said sites to ligate the fragment into CAG::OC1. A version of px458 that contains the EF1a promoter^17^ was modified to include CrxE1 immediately upstream of the EF1a sequence using Sal1 sites. To maintain the unique Bbs1 sites in the px458 vector for insertion of guide sequences, a Bbs1 restriction site in the CrxE1 sequence was mutated by one base pair (GTCTTC to GACTTC). The previously reported “Nrl guide 2”^12^ was then cloned into the base (EF1a) px458 vector as well as the modified CrxE1/px458 vector using Bbs1. All constructs generated through PCR were sequence verified.

### Electroporation

Ex vivo electroporation experiments were performed as previously reported^18^ with the following modifications. The electroporation chamber was rinsed with 70 uL of 1X Phosphate Buffered Saline (PBS) for a minimum of eight times between groups. Electroporation mixes were made using 50µL total volume in a 1X PBS final concentration, with plasmid DNA for each construct added to 10µg total (0.2µg/µL).

In vivo electroporations were performed as previously reported^1^ with the following modifications. Pulled glass needles (World Precision Instrument, 1B100F-4) were backfilled with each mix prior to electroporation. In vivo electroporation mixes were made using a total of 10µL volume in Tris-EDTA (TE), with each plasmid DNA added to 10µg total (1µg/µL). A Femtojet (Eppendorf) was used to inject DNA into the sub-retinal space. Tweezer-type electrodes (Bulldog Bio, CUY650P-10) were used to apply electrical pulses. A Nepagene NEPA21 Type II Super Electroporator (Bulldog Bio, NEPA21)was used for both ex vivo and in vivo electroporations.

### Ex vivo Explant Culture and Fixation

Retinas were cultured on 13 mm/0.2 micron floating filters (Cytiva, 10417001) on DMEM/F-12 media (Gibco, 11320082) with 10% Fetal Bovine Serum (Thermofisher, A3160602) and 1X L-Glutamine, Penicillin, Streptomycin (Gibco, 10378016). Retinas were fixed for 30 minutes at room temperature in 4% paraformaldehyde in 1XPBS.

### Immunofluorescence

Primary antibodies were diluted in a 1XPBS with 0.3% Triton X-100 solution. Primary antibodies used were rabbit anti-GFP 1:500 (ThermoFisher Scientific, A-6455), mouse anti-Rxrg 1:30 (Santa Cruz Biotechnology, SC-365252), mouse anti-Rhodopsin 1:500 (Santa Cruz Biotechnology, SC-57432), goat anti-Opn1SW 1:1000 (Santa Cruz Biotechnology, SC-14363), rabbit anti-RFP 1:250 (Rockland Immunochemicals, 600-401-379), and rabbit anti-Onecut1 1:300 (Proteintech, 25137-I-AP).

All secondary antibodies were obtained from Jackson Immunoresearch and were designated as appropriate for multiple labeling. Retinas were processed for staining as previously described^18^.

### Microscopy

All Confocal images were obtained using a Zeiss 800 confocal microscope. Images were analyzed using Image J and displayed using Affinity Designer software. All quantified images were analyzed blinded to the experimental condition. Files were duplicated and saved without identifiers, selected randomly, quantified, and then matched to their original file name.

## Results

### CrxE1 activity in postnatal mice is photoreceptor specific

To determine the activity of the CrxE1 element in the mouse retina in vivo, retinal electroporation was performed at p0 with a CrxE1::GFP reporter plasmid as well as a CAG::mCherry reporter plasmid (Figure 1). A minimum of 3 retinae, each from separate individuals, were dissected and fixed at p2, p8, and p14. All retinas were stained with 4’,6-Diamidino-2-Phenylindole (DAPI), and for retinas harvested at p8 and p14, a Rhodopsin antibody was used to visualize rod photoreceptor cells in the outer nuclear layer (ONL). The CrxE1 element was shown to drive activity specifically in the ONL, with only an occasional cell that appeared displaced into the inner nuclear layer (INL). At p8, 97.4 ± 2.2% of GFP+ cells were in the ONL, and at p14, 99.6 ± 0.6% of GFP+ cells were in the ONL (Supplemental Table 1). Analysis of sections by cell counting quantified an average of 9.02% CrxE1+ only cells out of total electroporated cells at p2, 14.46% at p8, and 14.12% at p14 (Figure1). Virtually every CrxE1+ only cell was positioned in the ONL and co-stained with Rhodopsin. Conversely, as the division between the INL and ONL becomes more apparent at P8 and P14, a significant portion of the CAG+ only cells were clearly located in the INL. Thus, targeting the CrxE1+ population in addition to the CAG+ population leads to increased reporting of photoreceptor cells.

**Figure 1.**
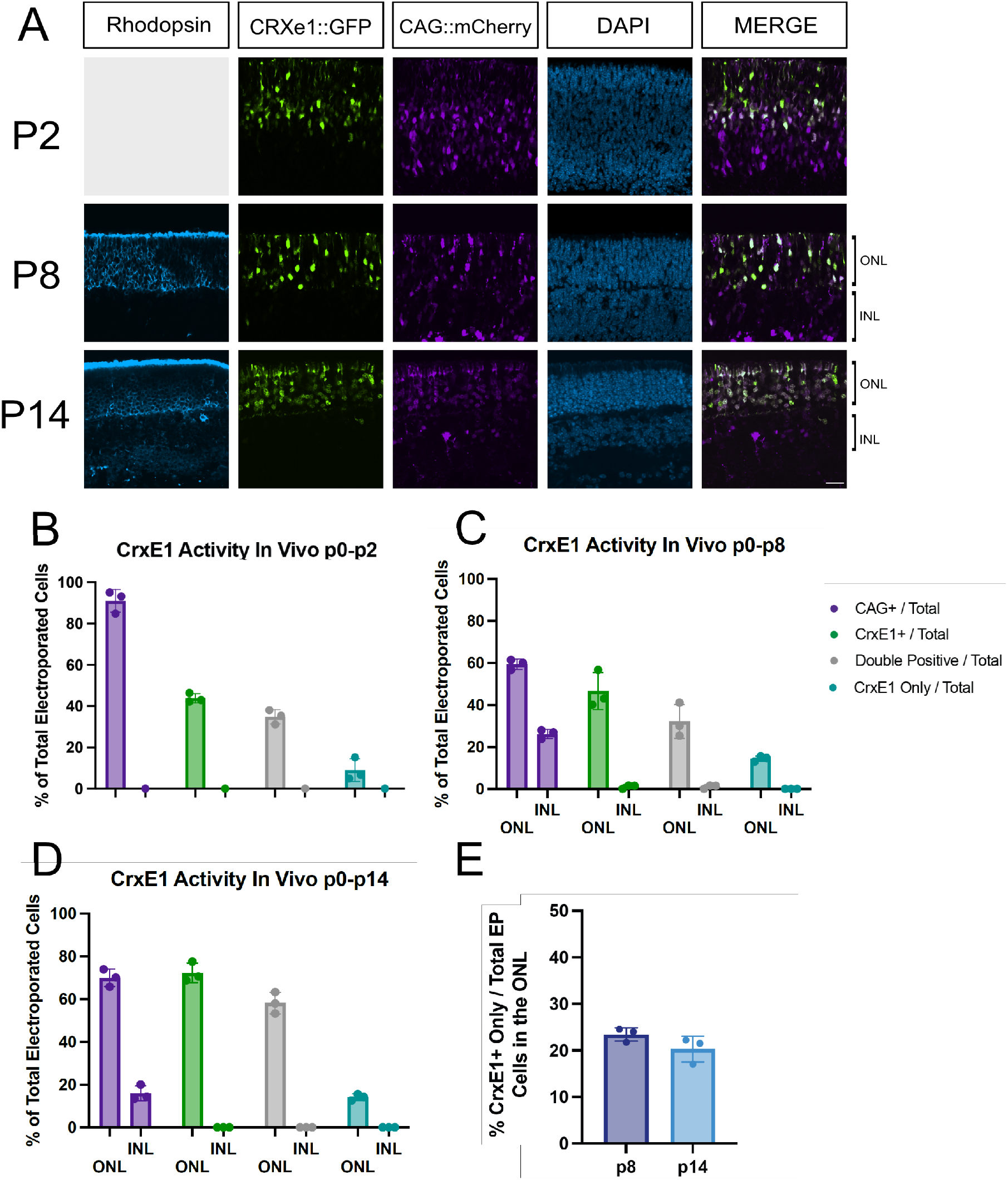
Characterization of CrxE1::GFP activity compared to CAG::mCherry activity across developmental timepoints. A. Confocal microscopy of retinal sections electroporated in vivo at p0 and processed at either p2, p8, or p14. Representative z-projections are shown with the developing ONL at the top of the section. Columns from left to right are Rhodopsin immunostaining in cyan, CrxE1::GFP epifluorescence, CAG::mCherry epifluorescence, DAPI staining in blue, and a merge of the CrxE1 and CAG reporter channels B-D. Quantification of reporter activity across 3 biological samples using total electroporated cells as the denominator and categorized for ONL or INL for panels C and D when the INL is apparent E. Quantification of CrxE1+ only cells out of electroporated cells specifically in the ONL as defined by Rhodopsin immunostaining. Scale bar in panel A is 20µm and applies to all panels. Abbreviations: ONL (Outer Nuclear Layer), INL (Inner Nuclear Layer), GCL (Ganglion Cell Layer), EP (Electroporated)

### Combining CrxE1 with a broadly active promoter increases reporter activity

The observation that as early as p2, the CrxE1 element is more strongly active than CAG in developing photoreceptors (Figure 1), led to the hypothesis that there is the potential to increase the number of reporter-positive cells within the electroporated population. We constructed a new single plasmid reporter which implemented both the widely used CAG element and the CrxE1 sequence to drive GFP. We tested multiple possible insertion points of CrxE1 into the CAG::GFP vector; one position located upstream of CAG, another between CAG and the GFP coding sequence, and an insertion point in the 3’UTR region of GFP. To screen for reporter activity, we co-electroporated p0 retinas ex vivo with one of the new GFP constructs and a control construct with CAG driving mCherry for p2 timepoints or TdT for p8 timepoints, and then cultured the retina for either 2 days or 8 days (Figure 2). It was immediately apparent that insertion of the CrxE1 sequence just 5’ of the GFP element in the plasmid did not increase activity, but actually led to the reporting of less cells than the original CAG::GFP vector. However, the other insertion points upstream of CAG or downstream of GFP increased the number of GFP+ cells proportional to the RFP+ cells. There was no consistent observable difference based on the orientation of the sequence in these positions. Insertion of the CrxE1 element just 5’ of the CAG element most consistently increased the number of cells targeted above the CAG alone condition, and by p8 we observe that GFP+ only cells localize in the ONL.

**Figure 2.**
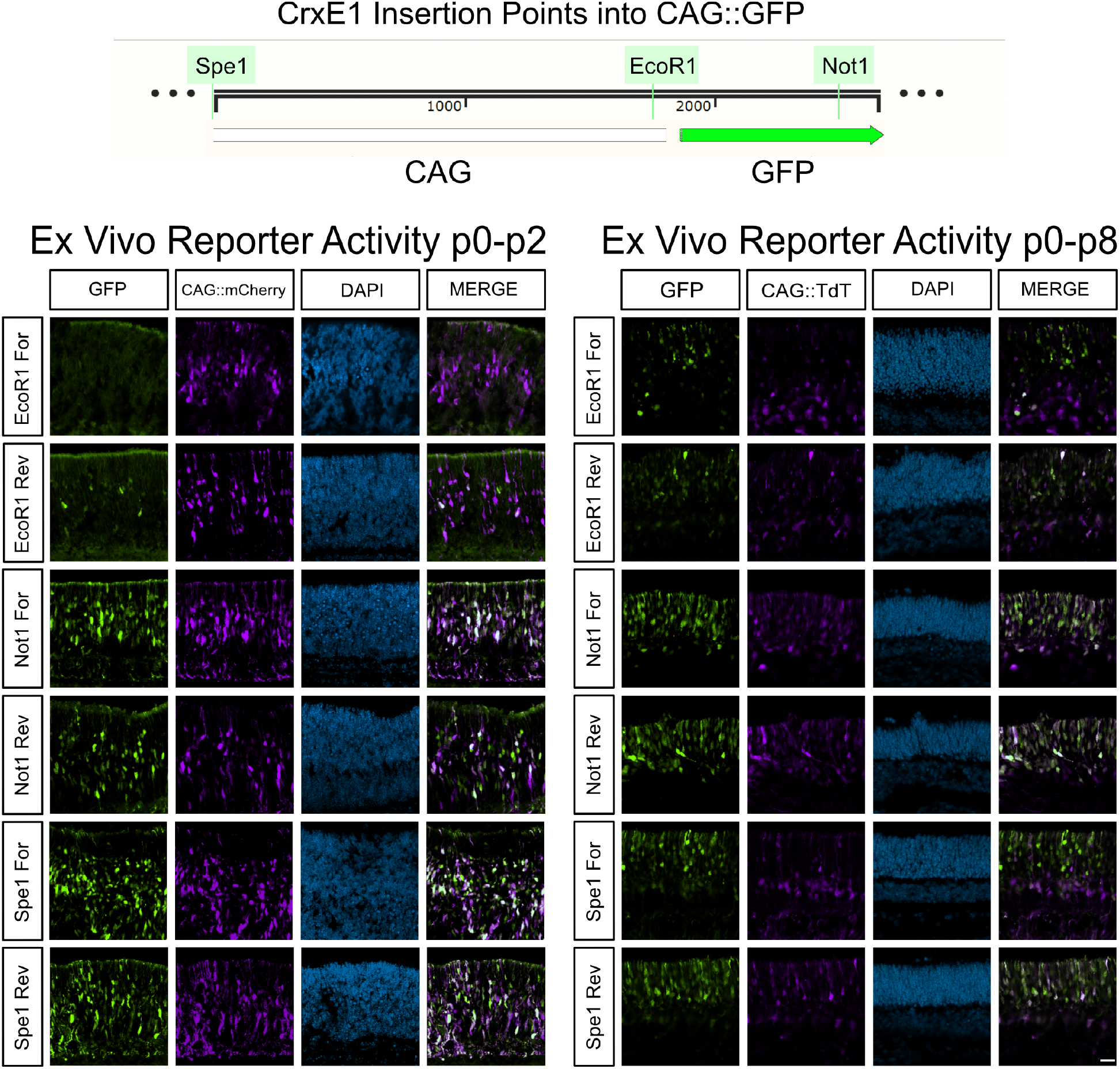
Differential CrxE1/CAG::GFP activity based on insertion point of CrxE1 into the CAG::GFP vector. Retinas were dissected and electroporated ex vivo and cultured for either 2 days or 8 days. Representative z-projections are shown with the developing ONL at the top of the section. Columns from left to right are GFP epifluorescence driven by various constructs, epifluorescence by either CAG::mCherry at p2 or CAG::TdT at p8, DAPI staining in blue, and a merge of the GFP and CAG reporter channels. Scale bar is 20µm and applies to all panels.

### The CrxE1/CAG element enhances the Onecut1 gain-of-function phenotype

After identification of the CrxE1/CAG::GFP reporter which showed an observable increase in the amount of photoreceptor cells targeted at p0, we next wanted to test whether a similar plasmid vector could be used to drive the expression of a gene of interest. This would provide us with a quantifiable indication of whether CrxE1 combined with CAG had an increased level of activity above CAG alone in the retina. We electroporated p0 mouse retinas in vivo with either a newly constructed CrxE1/CAG::OC1 plasmid or CAG::OC1, as well as our CrxE1/CAG::GFP reporter. Additionally, we included the ThrbCRM1::TdT reporter, which is active in RPCs that generate cones and is a direct target of the OC1 transcription factor. It has been reported that the ThrbCRM1 element is activated in the postnatal retina in response to Onecut1 overexpression^11^. We observed that as early as p14, an average of 40.36% of electroporated cells showed induced ThrbCRM1 activity in the CAG::OC1 group, whereas an average of 55.00% of electroporated cells showed induced ThrbCRM1 activity in the CrxE1/CAG::OC1 group (Figure 3). We performed a two tailed nested t-test analysis on the data generating this 14.6% ± 2.34% difference between groups and conclude a significant difference with a p-value of 0.0033. There was no significant difference between technical replicates across samples within a group (Supplemental Table 4).

**Figure 3.**
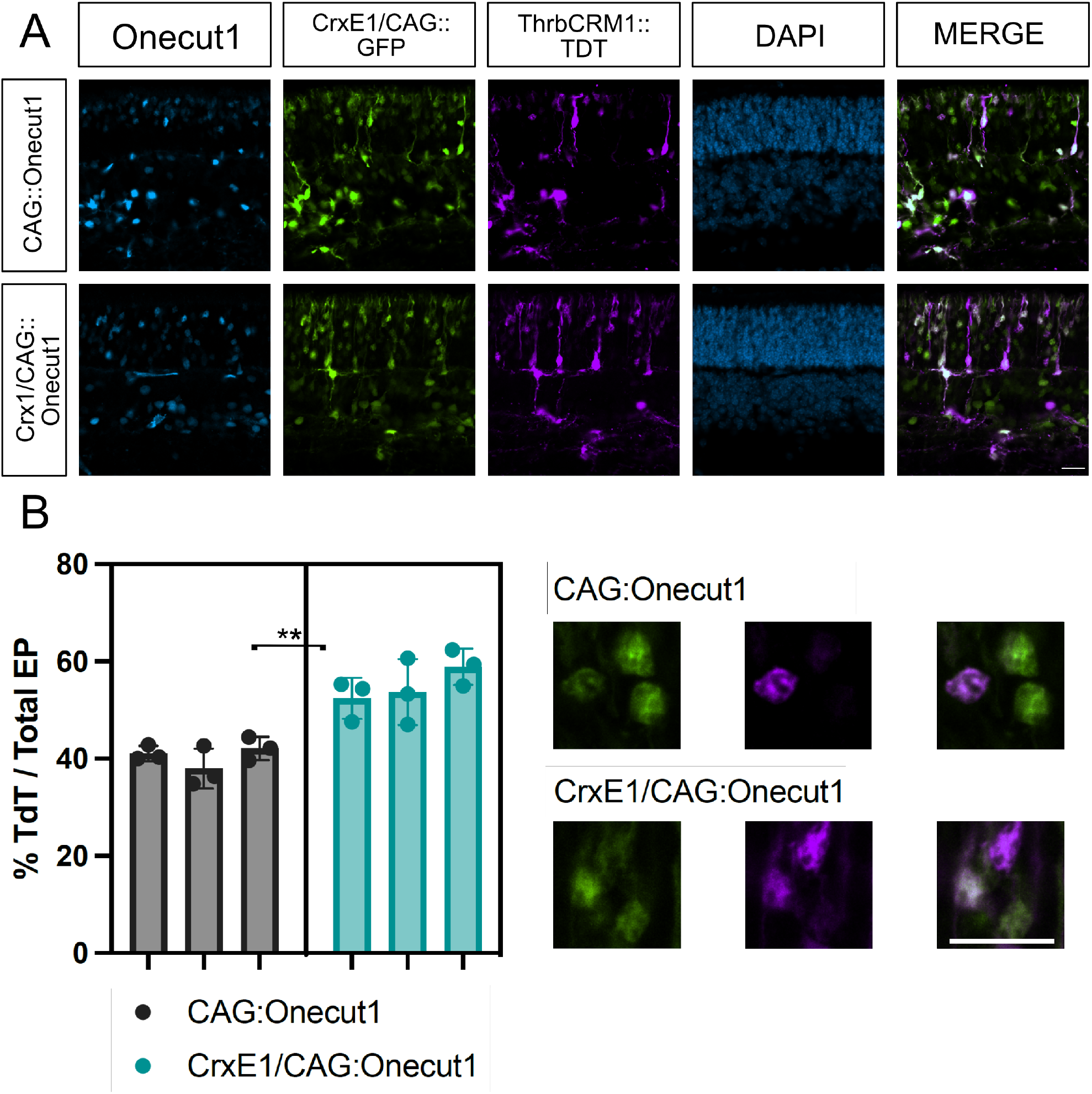
A. Confocal microscopy of retinal sections electroporated in vivo at p0 with CrxE1/CAG::GFP and either CAG::Onecut1 or CrxE1/CAG::Onecut1. Retinas were processed at p14 and sections were stained with a Onecut1 antibody. Representative z-projections are shown with the developing ONL at the top of the section. Columns from left to right are Onecut1 immunostaining in cyan, CrxE1/CAG::GFP epifluorescence, ThrbCRM1::TdT epifluorescence, DAPI staining in blue, and a merge of the GFP and TdT reporter channels B. Quantification of ThrbCRM1::TdT reporter and GFP double positive cells across 3 biological samples using CrxE1/CAG::GFP positive cells as the denominator. The statistical method used was the two tailed nested t-test analysis performed within the software program PRISM. A single asterisk indicates a p-value of p< 0.05, and a double asterisk indicates a p-value of p< 0.01. Scale bar in panel A is 20µm and applies to all panels.

### CrxE1 addition enhances a Cas9 loss-of-function phenotype

Finally, we wanted to test whether the addition of the CrxE1 sequence could enhance gene editing in the retina. We decided to test this in the context of the px458 vector system and guides which have been validated to work in the mouse retina^12^. Using a modified px458 vector with the ubiquitous promoter EF1a driving Cas9 and GFP, we inserted the CrxE1 element 5’ of the EF1a promoter to test whether it would increase the efficiency of gene editing in photoreceptors. To assess this, we focused on the Nrl transcription factor expressed by rod photoreceptors. In mice retinae with the px458 gene editing plasmid introduced in vivo at p0, rod cells show a upregulation of S-opsin at p21, consistent with the Nrl germline knockout phenotype^12^. We tested one of the Nrl targeting guide RNAs previously reported to be effective^12^. Retinas were electroporated with either the px458::Nrlg2 or CrxE1/px458::Nrlg2 plasmids in vivo at p0 and then analyzed at p21. Consistent with the original report using this guide, we observed that at p21 34.83% of all targeted cells activate S-opsin in response to the Nrl CRISPR condition in the px458::Nrlg2 group. By comparison, 49.47% of targeted cells activate S-opsin in the CrxE1/px458::Nrlg2 group (Figure 4). We performed a two tailed nested t-test analysis on this 14.6% ± 2.59% difference between groups and identified a significant difference with a p-value of 0.0049. There was no significant difference between technical replicates across samples within a group (Supplemental Table 5).

**Figure 4.**
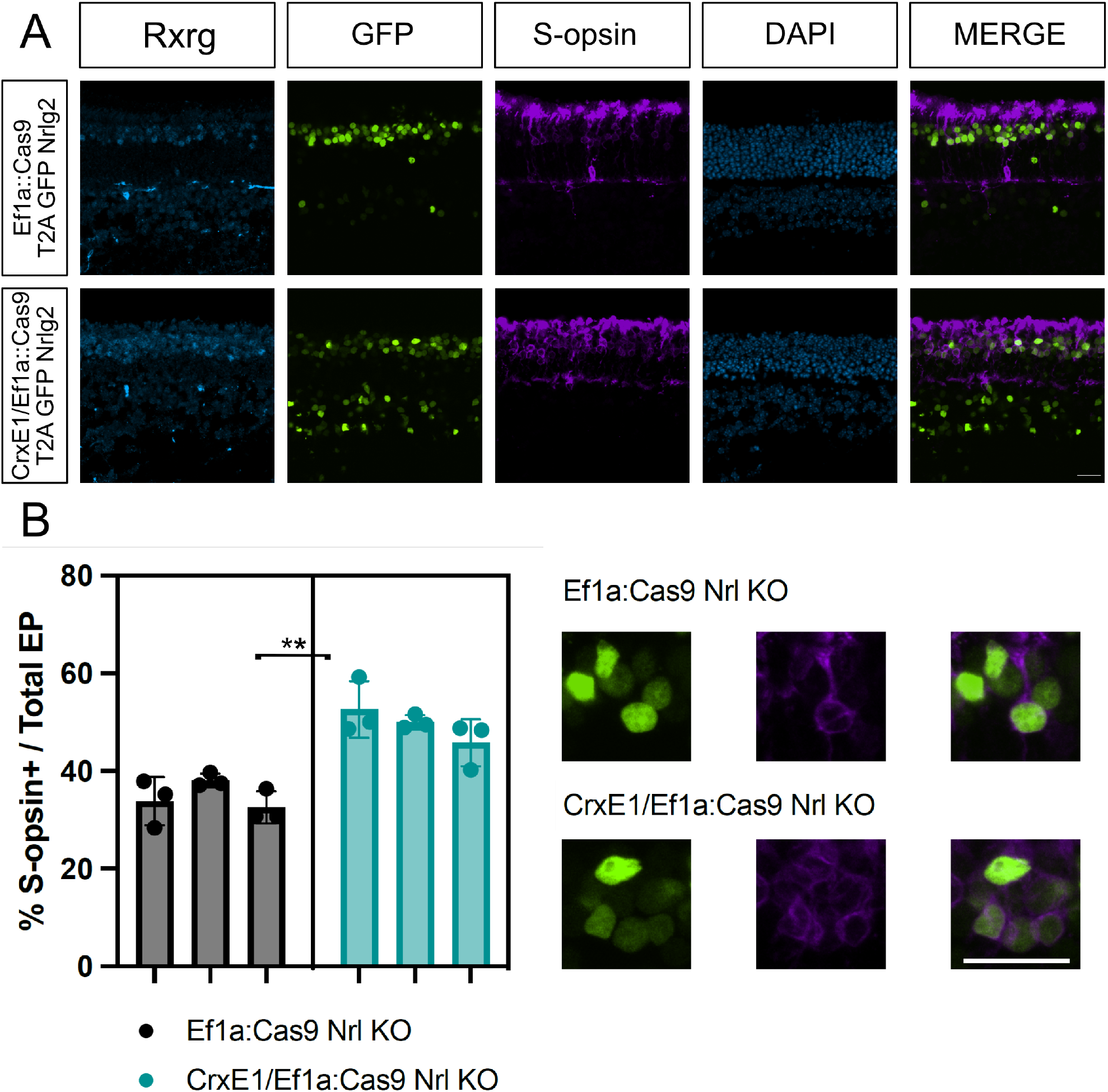
A. Confocal microscopy of retinal sections electroporated in vivo at p0 with either Ef1a::Cas9 T2A GFP Nrlg2 or CrxE1/Ef1a::Cas9 T2A GFP Nrlg2. Retinas were processed at p21 and sections were stained with a S-opsin antibody. Representative z-projections are shown with the developing ONL at the top of the section. Columns from left to right are Rxrg immunostaining in cyan, GFP epifluorescence, S-opsin immunostaining, DAPI staining in blue, and a merge of the GFP and TdT reporter channels B. Quantification of S-opsin and GFP double positive cells across 3 biological samples using GFP positive cells as the denominator. The statistical method used was the two tailed nested t-test analysis performed within the software program PRISM. A single asterisk indicates a p-value of p< 0.05, and a double asterisk indicates a p-value of p< 0.01. Scale bar in panel A is 20µm and applies to all panels.

## Discussion

CrxE1 activity was first predicted to be photoreceptor specific based on in silico studies using mouse DNA sequence^9^, and the sequence was shown to be active in this population. However, it was later characterized in chick and mouse retina tissue ex vivo, and while it showed the highest levels of activity in photoreceptors in both mouse and chick, it was also shown to be active in rod bipolar cells in mouse^10^. In the current study the degree of specificity of this element in the mouse retina was analyzed in vivo for the first time.

As early as p2 there is an observable difference between the localization of CrxE1 positive cells compared to cells reported on by the broadly active promoter CAG. The Crx gene is highly expressed in photoreceptor precursor cells, and this may indicate that the CrxE1 element is at least in part responsible for this early activity. While Crx gene expression is also seen to a lesser degree in bipolar cells, the in vivo activity of the CrxE1 element is highly specific to photoreceptors. By p8, rhodopsin protein expression delineates the outer nuclear layer (ONL) and at this developmental timepoint CrxE1 activity is shown to be specific to rod photoreceptors in the electroporated population. At this timepoint, we observed 23.9% of CAG::mCherry positive cells to be in the INL. Considering this proportion decreases throughout development, this is consistent with the approximately 20% of INL cells in the electroporated population which has been observed in the p10 rat retina^1^. By p14, the delineation between the INL and the ONL is most readily observable, and CrxE1 activity is shown to be consistently and exclusively in the ONL in mouse at this timepoint. The specific difference between the ex vivo activity of CrxE1 in some bipolar cells^10^ and the in vivo exclusion of activity in these cells is of note. The mechanism by which CrxE1 activity is silenced in bipolar cells may be specific to the in vivo environment.

Notably, there was a difference in the proportion of CrxE1+ cells to CAG+ cells as a function of timepoint, especially at p2. This is likely due to the fact that there are still multipotent RPCs targeted by CAG at p2, but by p8 this population has terminally divided to form postmitotic cells, which may or may not strongly activate CAG or the CrxE1 element depending on cell subtype. The higher standard deviation of this proportion seen at p2 may similarly be due to the position of the electroporated area relative to the center of the retina. Sections of the retina that are more peripheral will be lagging in developmental stage and potentially still have more RPCs.

Many of the cells targeted by the CrxE1 element can also be targeted by ubiquitous promoters, as shown in Figure 1. The strengths of using broadly active promoters include targeting a large number of cells, as well as intentionally driving gene expression in multiple cell subtypes. However, one negative impact of this approach is that the cells which are targeted may not be enriched for the cell subtype of interest. Importantly, CAG does not appear to be strongly active in newborn photoreceptors. As previously stated, Crx transcription begins very early in newborn photoreceptor cells, and we observe strong CrxE1 activity as early as p2 in a significant portion of cells where CAG is weakly active or not active at all.

The CrxE1 element, while highly specific to photoreceptors, is active in fewer cells than the CAG promoter, even in the ONL. Thus, it was important to determine whether a plasmid containing both a ubiquitous promoter and the CrxE1 element could increase the amount of photoreceptors that are normally targeted by CAG. To generate mRNAs utilizing CrxE1 in a plasmid, transcription must maintain an efficient start site, and must continue uninhibited through the intended open reading frame of a target sequence. Thus, the position and orientation of each promoter or enhancer sequence should be tested to determine efficient plasmid activity. Qualitative observations revealed clear differences in reporter activity based on the insertion points of CrxE1 into CAG::GFP (Figure 2). Despite the fact that promoters such as CAG are assumed to be broadly active in all cells, the results in this study indicate that incorporation of an additional cell subtype specific enhancer can increase overall activity of these elements.

The ability of Onecut1 overexpression at p0 in mouse to induce ThrbCRM1 activity by p14 is easily quantifiable using a previously described ThrbCRM1 reporter. Reporter activity of ThrbCRM1 when testing the CrxE1/CAG::OC1 plasmid increased proportionally with the number of cells targeted by CrxE1 alone (Figure 1, Figure 3). At p14, we observed an average of 14.1% CrxE1+ only cells across developmental timepoints (Supplemental Table 1). At p14, we also observe a 14.6% increase in ThrbCRM1::TdT cells when using the CrxE1 modified misexpression plasmid compared to the CAG::OC1 phenotype (Supplemental Table 2). This represents a 36% increase over baseline in electroporated cells that activate the ThrbCRM1 reporter. Similarly, in the loss-of-function experiments, we also observe a 14.6% increase in S-opsin positive electroporated cells at p21 when comparing the CrxE1 modified Cas9 plasmid to the Ef1a::Cas9 phenotype, which represents a 42% increase over baseline in the cells with a Nrl loss-of-function phenotype (Supplemental Table 3). These two sets of experiments employ different plasmid DNA backbone sequences, and one relies on Cas9 activity and sgRNA efficiency while the other does not. The similarity between the change in percent of phenotypic cells in this case is not due to an exceptionally low standard deviation in either set of experiments, but rather a uniquely common average of nine technical replicates across three biological samples for each. Nevertheless, we can conclude that the similar increase in phenotype is likely directly related to the number of cells that are targeted by the CrxE1 element in both settings.

It is therefore critical to point out that unlike the cells targeted by broadly active promoters, virtually every cell targeted by CrxE1 appears poised to respond to photoreceptor specific manipulations. This may be related to the early activity of CrxE1 in the context of cell birth. These cells which remain in the ONL throughout postnatal development are normally developing rod photoreceptors. However, these photoreceptors are at a stage in their development during which many of them can be manipulated starting at p0 to maintain a repression of rod programming, and instead produce cone associated proteins Rxrg and S-opsin^11,12^.

This increase in phenotypic penetrance is critical not only to this research but to any other experiments that target developing photoreceptors. Depending on the function of one’s gene of interest, this increase could strongly influence whether a novel phenotype is readily detectable or not. Additionally, there are genes which are expressed in both rod and bipolar cells, and targeting these genes using plasmids that do not contain any ubiquitous promoter, and instead with only the CrxE1 element, should be further investigated.

Importantly, one of the intended outcomes of this study was to interrogate the possibility of enhancing previously studied photoreceptor fate change phenotypes. These results directly apply to photoreceptor generation research and other translational research opportunities regarding treatment of blinding diseases. The reporter studies here within show that the CrxE1 element is strongly active in rod photoreceptor cells, including at an early stage in their development. Furthermore, the data reveal that conducting gain or loss-of-function experiments with this element in addition to a broadly active promoter can lead to changes in gene expression patterns that exceed what has been previously reported. The observed increase in the proportion of phenotypic cells re-emphasizes the importance of cell subtype specific targeting. This phenomena requires further study and will be the topic of future investigations.

## Acknowledgements

Acknowledgement is made to the donors of Macular Degeneration Research, a program of BrightFocus Foundation for support of this research (grant M2020157 to M.E.) and support by the National Eye Institute of the National Institutes of Health under award number R01EY024982 (to M.E.). The content is solely the responsibility of the authors and does not necessarily represent the official views of the BrightFocus Foundation or the National Institutes of Health. The authors thank members of the Emerson Lab for critically reading the manuscript.

